# Transcriptional coupling of telomeric retrotransposons with the cell cycle

**DOI:** 10.1101/2023.09.30.560321

**Authors:** Mengmeng Liu, Xiao-Jun Xie, Xiao Li, Xingjie Ren, Jasmine Sun, Zhen Lin, Rajitha-Udakara-Sampath Hemba-Waduge, Jun-Yuan Ji

## Abstract

Instead of employing telomerases to safeguard chromosome ends, dipteran species maintain their telomeres by transposition of telomeric-specific retrotransposons (TRs): in *Drosophila*, these are *HeT-A*, *TART*, and *TAHRE*. Previous studies have shown how these TRs create tandem repeats at chromosome ends, but the exact mechanism controlling TR transcription has remained unclear. Here we report the identification of multiple subunits of the transcription cofactor Mediator complex and transcriptional factors Scalloped (Sd, the TEAD homolog in flies) and E2F1-Dp as novel regulators of TR transcription and telomere length in *Drosophila*. Depletion of multiple Mediator subunits, Dp, or Sd increased TR expression and telomere length, while over-expressing E2F1-Dp or knocking down the E2F1 regulator Rbf1 (Retinoblastoma-family protein 1) stimulated TR transcription, with Mediator and Sd affecting TR expression through E2F1-Dp. The CUT&RUN analysis revealed direct binding of CDK8, Dp, and Sd to telomeric repeats. These findings highlight the essential role of the Mediator complex in maintaining telomere homeostasis by regulating TR transcription through E2F1-Dp and Sd, revealing the intricate coupling of TR transcription with the host cell-cycle machinery, thereby ensuring chromosome end protection and genomic stability during cell division.

## Introduction

Telomeres protect chromosome ends and play critical roles in chromosome replication and genome stability in species with linear chromosomes. While telomerase-based telomere maintenance is a common mechanism in most eukaryotes, some dipteran insects, such as *Drosophila melanogaster*, lack the telomerase enzyme and have evolved an exceptional strategy for telomere elongation. Apart from gaining a deeper understanding of the diversity, complexity, and richness of life on earth, studies of alternative strategies of telomere maintenance has helped elucidate the pressures and strategies taken by cancer cells to extend their replicative lifespan. In *Drosophila*, telomere elongation is accomplished through the transposition of telomeric-specific retrotransposons (TRs), which are specifically directed to chromosome ends and serve a critical role in maintaining chromosome stability. Decades of genetic and cytological studies have revealed that *Drosophila* utilizes three TRs – *HeT-A* (heterochromatic telomeric-associated sequence A), *TART* (telomere-associated retrotransposon), and *TAHRE* (Telomere-Associated and HeT-A-Related Element) – to extend tandem repeat sequences at the ends of its chromosomes (Abad et al., 2004; Levis et al., 1993; Traverse and Pardue, 1988). Following transcription by RNA polymerase II (Pol II), the sense-strand transcripts of *HeT-A* and *TART* are translated in cytoplasm to produce Gag proteins. These Gag proteins associate with the sense-strand RNAs from all the three TRs, transporting them back into the nucleus and to the chromosome ends, where they serve as templates for reverse transcription and incorporation at telomeres (Cacchione et al., 2020; Pardue and DeBaryshe, 2011; Pimpinelli, 2006). Translation and retrotransposition are constitutive processes, making transcriptional regulation of TRs, the initial step in their transposition life cycle, a primary determinant of telomere homeostasis.

Retrotransposons (Class I transposable elements) rely on Pol II dependent-transcription (Cordaux and Batzer, 2009; Wells and Feschotte, 2020). Activation of non-TR retrotransposons is associated with various health-related issues such as sterility, cancer, aging, and other diseases, and thus is regarded as detrimental (Cordaux and Batzer, 2009; Kazazian and Moran, 2017). Understanding how the TRs are regulated separately from all other retrotransposons is the key to understanding how dipteran cells have co-opted a cellular parasite to maintain its genome stability. In eukaryotes, the Mediator complex is the major co-activator of both protein-coding and most non-coding RNA gene transcription. Comprising approximately 30 distinct and conserved subunits, the Mediator complex is organized into four modules: the head, middle, tail, and CDK8 kinase modules (Allen and Taatjes, 2015). The head, middle, and tail modules can be purified together as the small core Mediator complex, which reversibly associates with the CDK8 kinase module (CKM) to form the large Mediator complex. This modular arrangement provides the Mediator complex with a large, flexible, and intricate surface, facilitating its interactions with a variety of transcriptional activators in diverse developmental and physiological contexts. Recent Cryo-EM studies have demonstrated that the Mediator head and middle modules directly interact with the Pol II C-terminal domain (CTD) (Schilbach et al., 2023), forming a molecular bridge between Pol II and enhancer-bound transcription factors (Allen and Taatjes, 2015).

Here, we report our analyses of the transcriptional regulation of *Drosophila* TRs, and present data illustrating how this regulation contributes to cell-cycle-dependent telomere homeostasis. We identified multiple subunits of the Mediator complex and two transcriptional factors, Scalloped (Sd) and E2F1-Dp, as novel regulators of TR transcription and telomere length in *Drosophila*. Through mutational analyses, we show that the large Mediator complex represses the transcription of TRs through Sd and E2F1-Dp, while the small Mediator complex is required for E2F1-Dp-dependent TR transcription. Additionally, CUT&RUN analyses reveal the direct binding of CDK8, Dp, and Sd to telomeric repeats. These results collectively illustrate the robust coupling of TRs with host cell-cycle machinery, jointly regulating telomere dynamics to ensure the success of the life cycle of TRs and the host cell genomic stability in *Drosophila*.

## Results

### Elevated telomeric-specific retrotransposon expression and increased telomere length in *Cdk8* and *CycC* mutants

Homozygous null mutants of Cdk8 (*Cdk8^K185^*) or CycC (*CycC^Y5^*) are lethal, but they survive to pupal stage due to maternal contributions of Cdk8 and CycC (Loncle et al., 2007; Xie et al., 2015). To understand the terminal defects in *Cdk8* and *CycC* mutants, we examined global gene expression profiles in late third-instar larvae using RNA-seq followed by “Pathway” and “Gene Ontology” cluster analyses (Boyle et al., 2004). As previously reported, we confirmed significant impact of *Cdk8* and *CycC* mutation on the transcription of genes involved in lipogenesis (via Sterol Regulatory Element-Binding Protein) (Li et al., 2022; Zhao et al., 2012), metamorphosis (via Ecdysone Receptor) (Xie et al., 2015), and DNA replication (via E2F1) (Morris et al., 2008; Zhao et al., 2013). Unexpectedly, we also observed a striking upregulation of certain non-coding RNAs, particularly retrotransposable elements. Among these, *HeT-A*, *TART*, and *TAHRE* are of particular interest due to their unique characteristics as telomeric retrotransposons (TRs), which play an essential role in the maintenance of telomere length in insects (Frydrychova et al., 2008; Mason et al., 2008; Mason, 2011; Pardue and DeBaryshe, 2011). We verified that these TRs were upregulated in both *Cdk8* and *CycC* mutants using qRT-PCR with the non-LTR retrotransposon *jockey* (*jockey gag*) and non-retrotransposon RNA Pol II-transcribed *RasGAP* gene as controls (Fig. 1A), confirming a novel role for Cdk8-CycC in negatively regulating the expression of TRs.

**Fig. 1.**
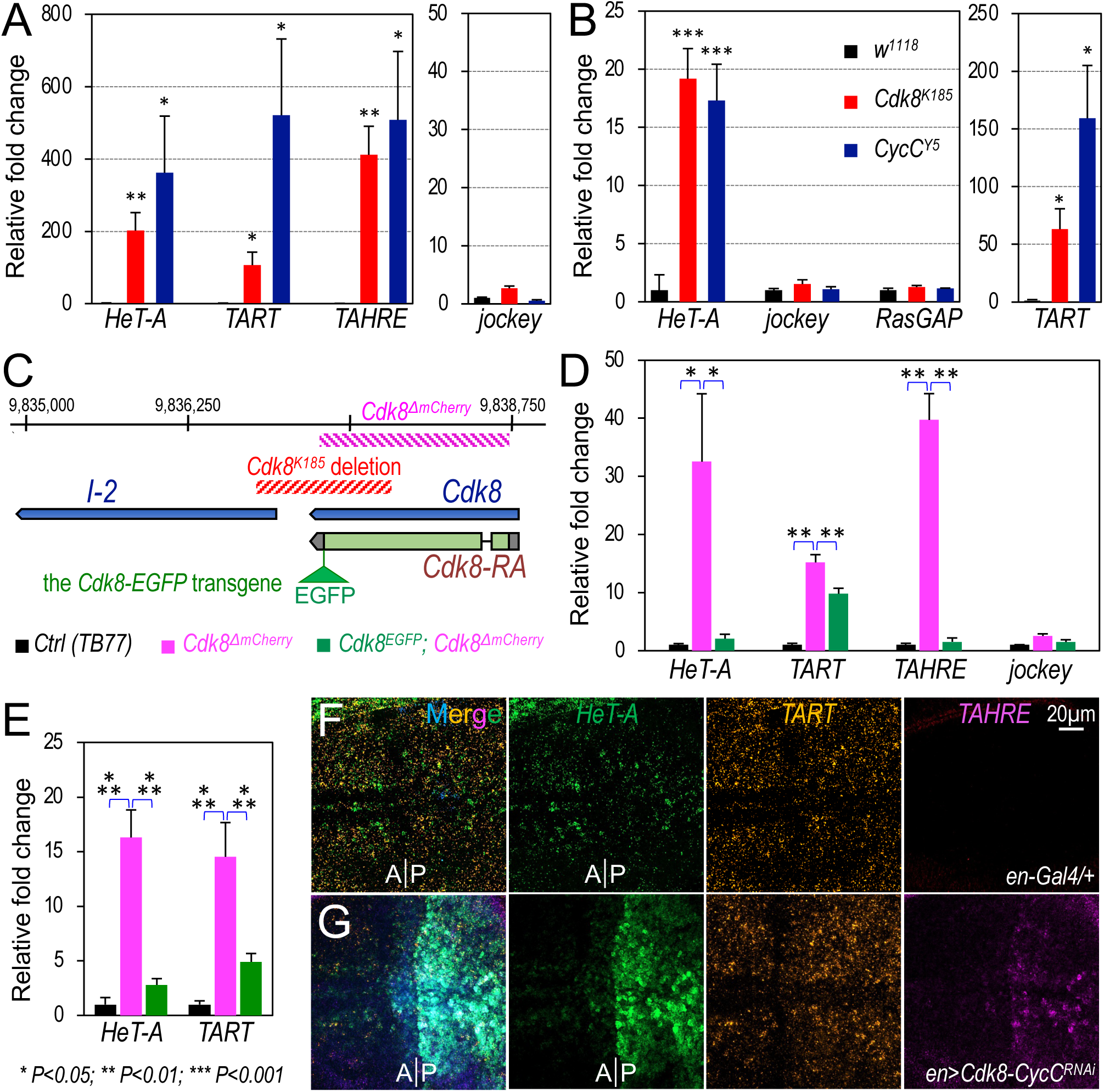
Increased expression of telomeric retrotransposons (TRs) in *Cdk8* and *CycC* mutants. (A) qRT-PCR analysis of *HeT-A*, *TART*, and *TAHRE* expression in *Cdk8^K185^*and *CycC^Y5^* mutants, with *w^1118^* serving as the control. The expression of *jockey gag* from the non-LTR-retrotransposon *jockey* (*jockey gag*) serves as a negative control. (B) Measurement of the telomere length by qPCR analysis of the three TRs using the genomic DNA from *Cdk8^K185^* and *CycC^Y5^* mutant larvae as the templates. *RasGAP* (the neighboring gene of *Cdk8*) and *jockey* serve as the specificity and negative controls. (C) Genomic region diagram of the *Cdk8* locus with deleted regions indicated by hatched lines. The EGFP tag is attached to the C-terminus of the Cdk8 protein, and the rescue construct is integrated into the second chromosome at the *attP40* site. (D) qRT-PCR analysis of the mRNA levels of *HeT-A*, *TART*, and *TAHRE* in *Cdk8^ΔmCherry^* (magenta), *Cdk8^EGFP^; Cdk8^ΔmCherry^* (green), and *TB77* (control, black). (E) Measurement of the telomere length by qPCR analysis using genomic DNA from *Cdk8^ΔmCherry^*, *Cdk8^EGFP^; Cdk8^ΔmCherry^*, or *TB77* larvae. (F/G) mRNA transcripts of *HeT-A* (green), *TART* (orange) and *TAHRE* (magenta) in the wing disc of *en-Gal4/+; UAS-BFP/+* (F, control) and *en-Gal4/+; UAS-Cdk8^RNAi^ CycC^RNAi^/UAS-BFP* (G) detected using the HCR RNA-FISH assay. * p<0.05, ** p<0.01, *** p<0.001 (one-tailed unpaired *t*-tests).

To evaluate the impact of deregulated TR expression in *Cdk8* and *CycC* mutants on telomere length, we used qPCR to measure telomere repeat copy number, an established method to quantify the TR repeats (Shpiz et al., 2011; Walter et al., 2007; Wylie et al., 2016), focusing on diploid larval brains to avoid the varying levels of ploidy in many other larval tissues. The copy numbers of *HeT-A* and *TART* were significantly increased (Fig. 1B), we confirmed this was due to increased telomere length by directly visualizing their length on polytene chromosomes. *Cdk8^K185^*, *CycC^Y5^*, or *Cdk8^ΔmCherry^*mutants (Fig. 1C) were independently crossed to *wild-type* flies to create heterozygotes of the mutations and to juxtapose homologous telomeres altered by the mutations with *wild-type* telomeres (Gao et al., 2010). Compared to the control (Fig. S1A), we observed uneven polytene chromosome ends in *Cdk8^K185^/+* (Fig. S1B), *CycC^Y5^/+* (Fig. S1C), and *Cdk8^ΔmCherry^*/+ larvae, showing that loss of *Cdk8* and *CycC* results in telomere elongation.

The three TRs at telomere form terminal repeats, called HTT (HeT-A, TART, and TAHRE) arrays (Frydrychova et al., 2008; Mason et al., 2008; Pardue and DeBaryshe, 2011), and proximal to each telomere’s HTT array are unique Telomere-Associated Sequences (TAS) comprising several kilobases of satellite DNA. Reporter genes inserted close to or within TAS are transcriptionally silenced. These telomeric position effect (TPE) have been used to investigate heterochromatin-induced gene silencing and telomere dynamics (Frydrychova et al., 2008; Mason et al., 2008; Pardue and DeBaryshe, 2011). Based on their roles in telomere length regulation, we predicted that *Cdk8* and *CycC* mutations would display dominant modification of TPE. We crossed the *Cdk8* and *CycC* mutant females to males bearing the fourth chromosome-linked TAS-inserted *118E-15* element, which places the *hsp70-white^+^* transgene under TPE (Fig. S1E) (Cryderman et al., 1999). Multiple mutant alleles, including *Cdk8^K185^* (Fig. S1F), *CycC^Y5^*(Fig. S1G), and *Cdk8^K185^CycC^Y5^* double mutants (Fig. S1H), strongly enhanced the TPE phenotype in males (Fig. S1E), further demonstrating novel roles of the Cdk8 Mediator module in telomere homeostasis in *Drosophila*.

### Validation of the role of Cdk8 and CycC in regulating telomeric-specific retrotransposon expression and telomere length

The original *Cdk8^K185^* null allele deletes part of *I-2* in addition to *Cdk8* (Fig. 1C) (Loncle et al., 2007; Xie et al., 2015), prompting us to generate a new *Cdk8* null allele using CRISPR-Cas9. We replaced the coding region of *Cdk8* with the *mCherry* gene to create the “*Cdk8^ΔmCherry^*” allele (Fig. 1C). Homozygous *Cdk8^ΔmCherry^* mutants were pupal lethal, we confirmed the lesion by sequencing and the lack of detectable Cdk8 protein (Fig. S1J). *Cdk8^ΔmCherry^*mutants were rescued to viability and fertility by a transgenic genomic fragment of the *Cdk8* locus with an EGFP tag at the C-terminus of Cdk8 (“*Cdk8^+^-EGFP*”) (Xie et al., 2015).

We tested whether the expression of TRs is altered in the *Cdk8^ΔmCherry^*mutant larvae. The expression of TR transcripts is significantly higher in *Cdk8^ΔmCherry^*mutant larvae than in the rescuing *w^1118^; Cdk8^+^-EGFP; Cdk8^ΔmCherry^* and *wild-type TB77* control larvae (Fig. 1D). In contrast, little difference was detected with the levels of the non-telomeric retrotransposon *jockey* (Fig. 1D). Telomere length in *Cdk8^ΔmCherry^*mutants was also significantly increased compared to controls (Fig. 1E). There was a detectable and statistically robust difference between the telomere lengths of *TB77* and *w^1118^; Cdk8^+^-EGFP; Cdk8^ΔmCherry^*, perhaps reflecting a gradual shortening in fly strains that had been mutant until the rescuing transgene was introduced. The longer telomeres in *Cdk8^ΔmCherry^* mutants were confirmed by direct visualization in polytene chromosomes from *Cdk8^ΔmCherry^/+* heterozygotes (Fig. S1D). Consistent with these observations, the *Cdk8^ΔmCherry^*allele also enhanced the telomeric position effects (Fig. S1I).

To test whether the effects of Cdk8 and CycC mutation on TR expression is chronic or acute, we utilized hybridization chain reaction RNA fluorescence *in situ* hybridization (HCR RNA-FISH), a sensitive technique that allows multiplexed quantitative detection of mRNA transcripts in individual cells (Choi et al., 2018). Endogenous *HeT-A* is known to be predominantly expressed in replicating diploid tissues such as imaginal discs and the larval neuroblasts (George and Pardue, 2003; Walter and Biessmann, 2004; Zhang et al., 2014). In wing discs, low levels of *HeT-A* and *TART* are detected in the wing pouch area, while the *TAHRE* transcripts are barely detectable (Fig. 1F). When Cdk8 and CycC were depleted in the posterior compartment of wing discs using *en-Gal4* driven RNAi, the expression of all three TRs was elevated in the cells of the posterior compartment compared to those in the anterior compartment (Fig. 1G). Compared to the control (Fig. S1K), significantly increased expression of TRs was also observed in polyploid salivary gland cells from *Cdk8^ΔmCherry^* mutant larvae (Fig. S1L). These findings show that depletion of Cdk8-CycC acts acutely to stimulate the expression of TRs in both polyploid and diploid cells.

### Mutations of additional Mediator subunits result in elevated telomeric-specific retrotransposon expression and increased telomere length

Cdk8 and CycC are known subunits of the Mediator complex. We investigated whether mutations of other Mediator subunits also altered TR expression and telomere length. MED7 is a subunit of the “middle” module of the Mediator complex and facilitates the assembly of the Mediator-Pol II holoenzyme (Sato et al., 2016). The uncharacterized *dMed7^MI10755^* allele harbors a Minos-based mutagenic gene trap cassette inserted in the 3’UTR region of *dMed7* locus (Fig. S2C), which we validated by PCR of genomic DNA (Fig. S2B). Homozygous *dMed7^MI10755^*mutants survive until third instar (Fig. S2C). In those mutants, we observed that expression of *HeT-A* and *TART* were significantly increased (Fig. 2A, Fig. S2D), and expression correlated with a significant increase of telomere length quantified using qPCR (Fig. 2B). Using HCR RNA-FISH, we observed that depletion of dMed7 specifically in salivary gland cells significantly increased the expression of *HeT-A* and *TART* (Fig. 2D) compared to controls (Fig. 2C). We also found that *Med7^MI10755^* dominantly enhanced telomeric position effects (Fig. S2E).

**Fig. 2.**
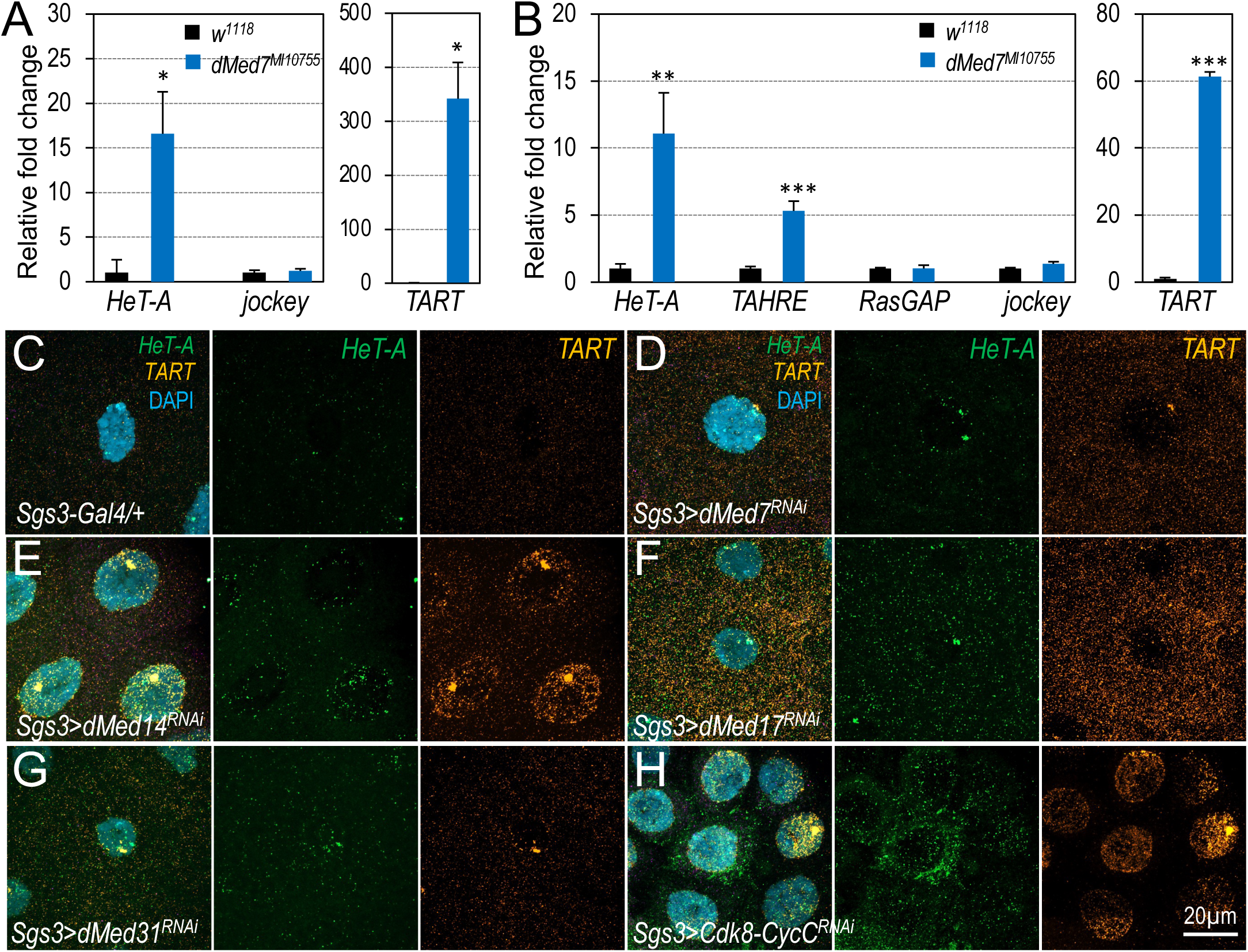
Disrupting additional Mediator subunits leads to elevated expression of TRs. (A) qRT-PCR analysis of *HeT-A* and *TART* expression in *dMed7^MI10755^* mutant larvae. (B) Measurement of the telomere length by qPCR analysis of TRs using genomic DNA from *dMed7^MI10755^* mutant larvae. * p<0.05, ** p<0.01, *** p<0.001 (one-tailed unpaired *t*-tests). (C-H) mRNA transcripts of *HeT-A* (green) and *TART* (orange) detected using the HCR RNA-FISH technique in salivary glands of indicated genotypes. Detailed genotypes: (C) *Sgs3-Gal4/+;+* (control); (D) *Sgs3-Gal4/+; UAS-dMed7^RNAi^/+*; (E) *Sgs3-Gal4/+; UAS-dMed14^RNAi^/+*; (F) and *Sgs3-Gal4/+; UAS-dMed14^RNAi^/+*; (G) *Sgs3-Gal4/+;UAS-dMed31^RNAi^/+*; and (H) *Sgs3-Gal4/+; UAS-Cdk8^RNAi^ CycC^RNAi^/+*.

Of the 30 Mediator complex subunits, MED14 and MED17 are essential central scaffold subunits for assembling the small Mediator complex (Chen et al., 2021). To test whether the small Mediator complex is also involved in regulating TR transcription, we depleted them in salivary gland cells and detected increased transcription of *HeT-A* and *TART*. Similar to dMed7, depletion of dMed14 (Fig. 2E), dMed17 (Fig. 2F), dMed31 (Fig. 2G), and Cdk8 and CycC (Fig. 2H) in salivary gland cells also resulted in an up-regulation of *HeT-A* and *TART* expression. Therefore, depleting multiple key Mediator subunits led to similar effects as the loss of Cdk8 and CycC, resulting in elevated transcription of TR in different types of cells. The most parsimonious interpretation of these observations is that the large Mediator complex acts as a transcriptional repressor of TRs.

### Increased telomeric-specific retrotransposon expression and telomere length in *scalloped* mutants

Despite decades of research into the mechanisms of TRs in regulating telomere length in *Drosophila*, the specific transcription factors that control TR transcription remain unknown. In the budding yeast *Saccharomyces cerevisiae*, two transcriptional activators – Ste12 and Tec1 – are reported to regulate the expression of the Ty1 retrotransposons (Aristizabal et al., 2015). In addition, certain non-essential yeast Mediator subunits regulate the expression of Ty1 (Salinero et al., 2018). Our BLAST search identified Scalloped (Sd) as a potential Tec1 homologue in *Drosophila*; we could find no Ste12 homologues. Sd is a TEAD/TEF-family transcription factor that acts downstream of the Hippo signaling pathway, which plays a crucial and conserved role in regulation of organ size. Hippo signaling is often found to be dysregulated in various human cancers (Halder and Johnson, 2011; Hansen et al., 2015; Pan, 2010; Wu et al., 2008; Yu et al., 2015; Zhang et al., 2008). Retrotransposons are a key source of genetic variation, and their abnormal expression and insertions have been observed in cancer (Carreira et al., 2014; Rodic et al., 2015), which prompted us to investigate whether Sd plays a role in TR expression or telomere length maintenance.

To test whether the expression of TRs and telomere length are affected in *sd^1^* mutants, we conducted similar qRT-PCR and qPCR assays. *sd^1^* is a hypomorphic allele induced by X-ray irradiation. *sd^1^* homozygotes are fully viable, but display a notched wing phenotype (Paumard-Rigal et al., 1998). We found that the mRNA levels of all three TRs are significantly elevated in *sd^1^* mutant larvae and dissected larval brain (Fig. 3A). Consistent with this, TR copy number was significantly higher in *sd^1^* mutant larvae when compared to *jockey* (Fig. 3B). Copy number was also higher in *sd^1^* mutants compared to mutants in *p53* (*Drosophila TP53*; Fig. 3B), which represses the expression of retrotransposons in *Drosophila,* zebrafish, and mammalian cells (Wylie et al., 2016). The longer telomeres were visualized in polytene chromosomes of *sd^1^/+* larvae (Fig. S3A). As with Mediator subunits, *sd^1^* also acted as a strong enhancer of TPE (Fig. S3B). Collectively, these observations indicate that Sd may be responsible for recruiting Mediator to telomeres, thereby repressing their expression.

**Fig. 3.**
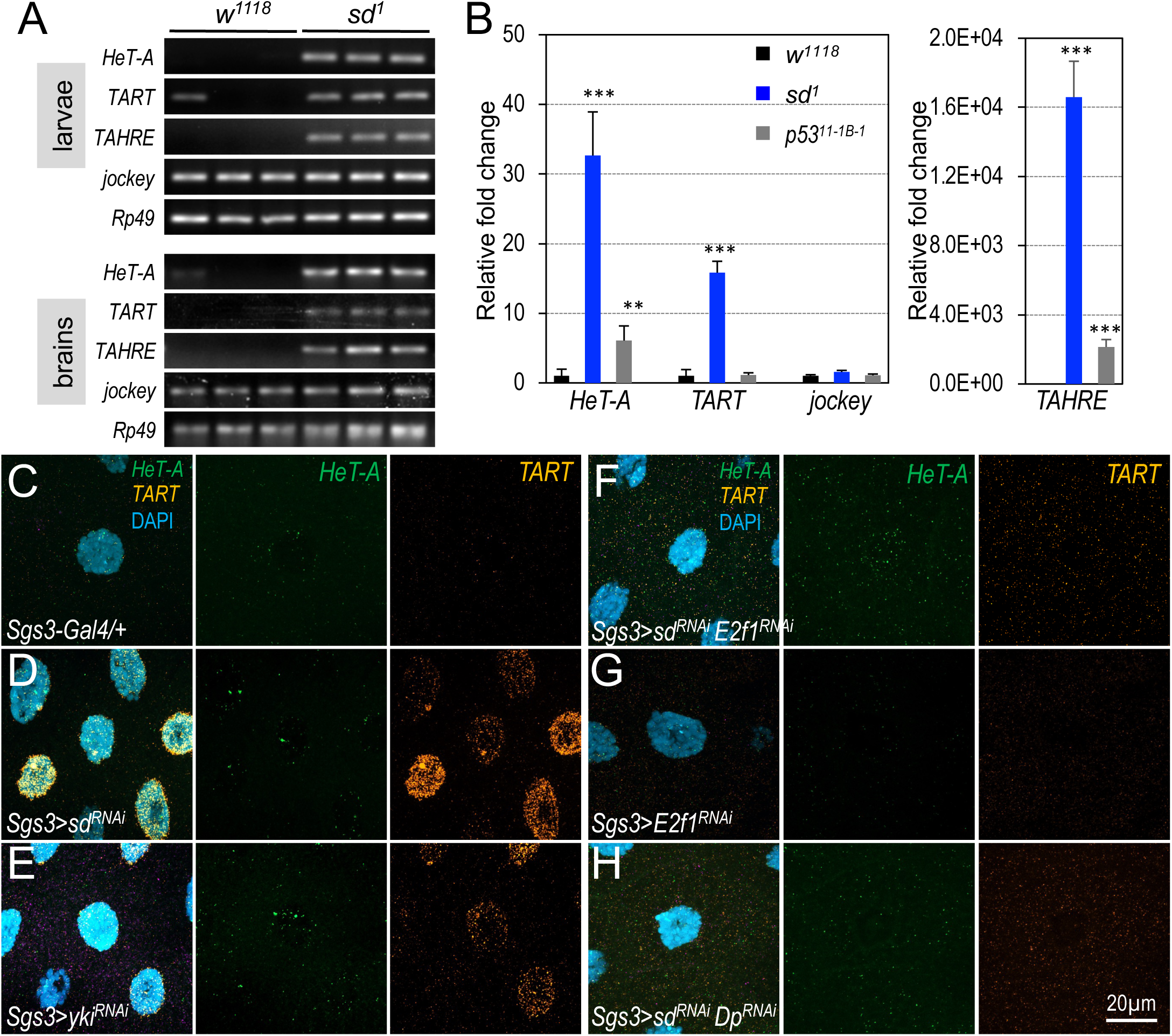
Impacts of Sd and Yki mutation on TR expression and telomere length. (A) qRT-PCR analysis of the expression levels of TR transcripts in both *sd^1^* homozygous mutant larvae and larval brain samples, and the results were visualized through nucleic acid gel electrophoresis. (B) Measurement of the telomere length by qPCR analysis using genomic DNA from *w^1118^*(control), *sd^1^* mutants, and *p53^11-1B-1^* mutant larvae. ** p<0.01, *** p<0.001 (one-tailed unpaired *t*-tests). (C-H) mRNA transcripts of *HeT-A* (green) and *TART* (orange) detected using the HCR RNA-FISH assay in salivary glands. Detailed genotypes: (C) *Sgs3-Gal4/+;+* (control); (D) *Sgs3-Gal4/+; UAS-sd^RNAi^/+*; (E) *Sgs3-Gal4/+; UAS-yki^RNAi^/+*; (F) *Sgs3-Gal4/+; UAS-sd1^RNAi^/UAS-E2f1^RNAi^*; (G) *Sgs3-Gal4/+;UAS-E2f1^RNAi^/+*; and (H) *UAS-sd1^RNAi^/UAS-Dp^RNAi^*.

Yorki (Yki) physically and genetically interacts with Sd (Goulev et al., 2008; Wu et al., 2008; Zhang et al., 2008). To assess the roles of Sd and Yki in TR expression, we depleted them individually from salivary glands and then examined TR expression using HCR RNA-FISH. Depleting either *sd* or *yki* significantly increased the levels of *TART*, though the effects on *HeT-A* and *TAHRE* were less pronounced (Fig. 3D and 3E) compared to controls (Fig. 3C). Similarly, depletion of *sd* in the posterior compartment diploid cells of wing discs using *en-Gal4* also resulted in elevated levels of *HeT-A* and *TART* (Fig. S3C), suggesting a negative regulatory role of both *sd* and *yki* in the transcription of *HeT-A* and *TART*.

The effects on TR expression resulting from depletion of *sd* and *yki* closely resemble the phenotypes observed upon depletion of multiple subunits of the Mediator complex. Mass spectrometry analyses of both *Drosophila* and cultured human cancer cells indicate that Yki and its human homolog YAP bind multiple Mediator subunits including Med1, Med12, Med14, Med15, Med19, Med23, Med24, and Med31 (Galli et al., 2015; Oh et al., 2013). We therefore speculate that the Mediator complex is recruited to the telomeric DNA through a Sd-Yki complex, which together repress the expression of the TRs.

### E2F1-Dp regulates the transcription of telomeric-specific retrotransposons and telomere length

Considering that a loss of transcriptional repression does not necessitate transcriptional activation, we investigated whether the transcription of TRs relies on any specific transcriptional activators. We considered E2F1 and its dimerization partner Dp as likely regulators of TR expression for three reasons. First, TRs are primarily expressed in proliferating cells, predominantly expressed in early S phase (Zhang et al., 2014). Notably, E2F1-Dp and other components of the Rb-E2F pathway are crucial regulators that control the transcription of genes involved in DNA replication during the transition from G1 phase to S phase of the cell cycle (Dimova and Dyson, 2005). Second, Cdk8 interacts with E2F1 and inhibits E2F1-dependent gene expression in *Drosophila* (Morris et al., 2008). Third, E2F1 can compete with Yki for binding to Sd, forming a E2F1-Sd repressor complex that regulates cell survival and organ size (Zhang et al., 2017).

To investigate the potential role of the Rb-E2F regulatory network in telomere biology, we first test determined whether ectopic expression of E2F1-Dp could induce the expression of the TRs. Using HCR RNA-FISH, we observed a significant increase in the expression of *HeT-A* and *TART* when E2F1 and Dp were co-expressed in salivary gland cells (Fig. 4A), suggesting the sufficiency of E2F1-Dp in the expression of TRs. During G1 phase of the cell cycle, Rbf1, the Rb homolog in *Drosophila*, acts as a repressor of E2F1-Dp activity (Dimova and Dyson, 2005). We therefore predicted that depletion of Rbf1 would also stimulate the transcription of TRs. Indeed, the depletion of Rbf1 resulted in a significant induction of *TART* expression and, to a lesser extent, the expression of *HeT-A* (Fig. 4C). In wing disc cells, ectopic expression of E2F1-Dp or depletion of Rbf1 also resulted in a strong up-regulation of all three TRs (Fig. S4).

**Fig. 4.**
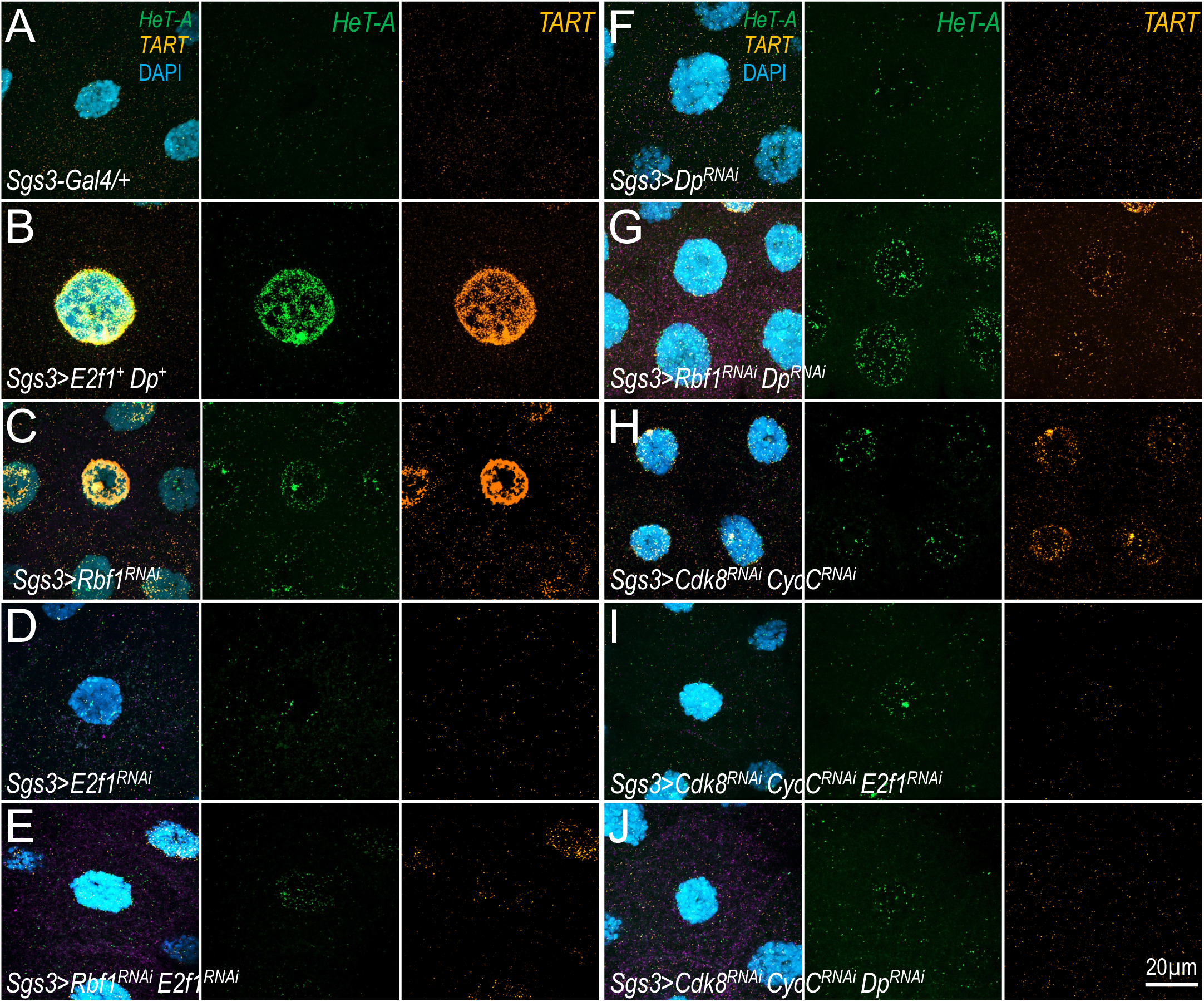
The role of RBF1-E2F1-Dp complex in regulating TR expression. mRNA transcripts of *HeT-A* (green) and *Tart* (orange) in salivary glands were detected using the HCR RNA-FISH assay. Detailed genotypes: (A) *Sgs3-Gal4/+;+* (control); (B) *Sgs3-Gal4/+; UAS-E2f1^+^ UAS-Dp^+^/+*; (C) *Sgs3-Gal4/+; UAS-Rbf1^RNAi^/+*; (D) *Sgs3-Gal4/+; UAS-E2f1^RNAi^/+*; (E) *Sgs3-Gal4/+; UAS-Rbf1^RNAi^/UAS-E2f1^RNAi^*; (F) *Sgs3-Gal4/+; UAS-Dp^RNAi^/+*; (G) *Sgs3-Gal4/+; UAS-Rbf1^RNAi^/UAS-Dp^RNAi^*; (H) *Sgs3-Gal4/+; UAS-Cdk8^RNAi^ CycC^RNAi^/+*; (I) *Sgs3-Gal4/+; UAS-Cdk8^RNAi^ CycC^RNAi^/UAS-E2f1^RNAi^*; and (J) *Sgs3-Gal4/+; UAS-Cdk8^RNAi^ CycC^RNAi^/UAS-Dp^RNAi^*.

We also predicted that the elevated expression of TRs upon *Rbf1* loss would be dependent on E2F1-Dp. Compared to wild-type larval salivary gland cells (Fig. 4A), co-depletion of Rbf1 and either *E2F1* (Fig. 4E) or *Dp* (Fig. 4G) strongly suppressed the effects of Rbf1 depletion (Fig. 4C) on *HeT-A* and *TART* expression. These observations are consistent to the model that Rbf1 inhibits E2F1-Dp-dependent gene transcription (Dimova and Dyson, 2005). We expanded this epistasis analysis by investigating whether the effects of Sd or Cdk8-CycC depletion were also dependent on E2F1-Dp. Compared to controls that deplete *sd* alone (Fig. 3D), co-depletion of *sd* with either *E2F1* (Fig. 3F) or *Dp* (Fig. 3H) significantly suppressed the ectopic expression of *HeT-A* and *TART*. Similarly, co-depletion of *Cdk8-CycC* with either *E2F1* (Fig. 4I) or *Dp* (Fig. 4J) also strongly reduced the effects of Cdk8-CycC knockdowns (Fig. 4H). These observations show that the effects of disrupted Mediator complex or Sd on TR expression are dependent on E2F1-Dp, strongly suggesting that the Mediator complex modulates telomere length homeostasis by restraining the expression of TRs through the transcription factors Sd and E2F1-Dp.

### Cdk8, Dp, and Sd directly bind to the telomeric HTT repeats

Our model predicted direct binding of the Mediator complex, E2F1-Dp, and Sd to the HTT repeats. To investigate this prediction, we utilized the CUT&RUN (Cleavage Under Targets & Release Using Nuclease), a sensitive high-throughput method to map genomic binding of chromatin-associated proteins (Skene and Henikoff, 2017). One notable technical challenge was the lack of “ChIP-grade” antibodies specific to Cdk8, E2F1-Dp, and Sd. Instead, for Cdk8 we used CRISPR-Cas9 to introduce an EGFP tag to the endogenous *Cdk8* gene (see Materials and Methods). For Sd and Dp, we used two existing EGFP-tagged lines where the endogenous genes were tagged with EGFP. Importantly, all three strains – *Cdk8^EGFP^*, *Dp^EGFP^*, and *Sd^EGFP^* – are fully viable and fertile, suggesting that the introduction of EGFP tags does not disrupt the normal functions of these three proteins.

Using wing discs from these EGFP-tagged lines, we investigated the genome-wide binding profiles of Cdk8, Dp, and Sd by CUT&RUN. We investigated binding at the telomeres of the long left arm of the *X* chromosome (*XL*) and the right arm of chromosome *4* (*4R*), which have been assembled by the *Drosophila* Heterochromatin Genome project (George et al., 2006). Our analyses of the assembled XL telomere region, spanning approximately 130 kilobases, revealed approximately 20 called peaks of Cdk8 binding, approximately called peaks of Dp binding, and two called peaks of Sd binding. Although the number of called peaks for Sd is lower than that of Cdk8 and Dp, a significant portion of binding peaks exhibit overlap among these three proteins (Fig. 5A). In fact, several “weaker” not-called Sd peaks overlapped with common Cdk8- and Dp-peaks, suggesting that all three may be present at many sites.

**Fig. 5.**
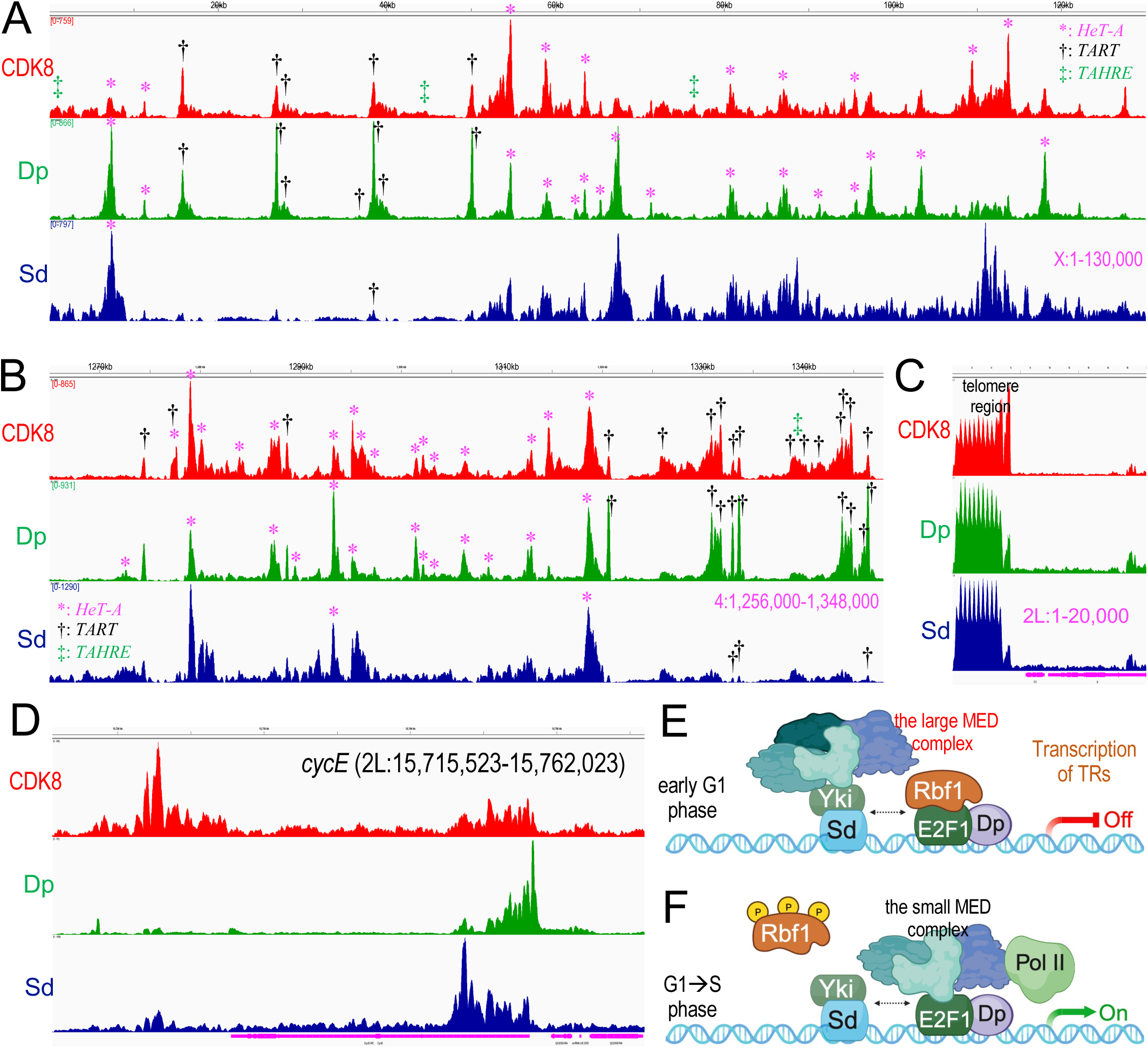
Direct binding of Cdk8, Dp, and Sd to telomeric HTT repeats. Genomic tracks display binding peaks for Cdk8 (red), Dp (green), and Sd (blue) at the telomeres of four different chromosome regions: XL (A, with the chromosome end on the *left*), 4R (B, chromosome end on the *right*), 2L (C), and *cycE* locus (D). To identify the specific sequences within the XL (A) and 4R (B) telomeres, we conducted individual BLAST searches and the locations of three TRs were marked using different symbols (* represents *HeT-A*, † represents *TART*, and ‡ represents *TAHRE*). (E/F) A model for the transcriptional regulation of TRs: during the early G1 phase, the large Mediator complex represses TR transcription via Sd and E2F1-Dp (E). As the cell progresses through the G1-S phase transition, the small Mediator complex becomes essential for activating TR transcription, primarily through E2F1-Dp (F).

Because these telomeric regions are not annotated in the *Drosophila* genome assembly Release r5.9, we conducted BLAST searches using the sequences associated with each of the called peaks. Our results revealed a significant overlap between most of these peaks and the three TRs, *HeT-A*, *TART* and *TAHRE* (Fig. 5A), supporting our model that Cdk8, Dp, and Sd directly regulate the transcription of TRs. Similarly, our analyses of the telomere of *4R* spanning approximately 92 kilobases revealed over 30 called peaks of Cdk8, over 20 called peaks of Dp, and five called peaks of Sd. Again, we observed substantial overlap (Fig. 5B). As with the telomere of *XL*, all of these called peaks also correlated with the three TRs (Fig. 5B). Notably, multiple *HeT-A* elements or *TART* elements cluster at the telomeres of XL and 4R; similar *HeT-A* or *TART* clustering pattern was depicted previously (George et al., 2006). As a positive control, we confirmed distinct binding patterns of Cdk8, Dp and Sd on their common target gene *cyclin E* (Fig. 5D). Binding of these three proteins were also found in the telomere regions of the chromosome 2L (Fig. 5D) and 3L (Fig. S5C), but those binding profiles could not be further analyzed without an assembly of the repetitive DNA at these telomeres. For a similar reason, we could not map CUT&RUN sequencing data for telomeres of *2R* (Fig. S5A), *3R* (Fig. S5D), and *4L* (Fig. S5B). Nevertheless, these observations provided further compelling evidence supporting the direct regulation of TR transcription by Cdk8, Dp (presumably together with its partner E2F1), and Sd.

## Discussion

Telomeres protect chromosome ends and play critical roles in chromosome replication and genome stability. Our results show that the large Mediator complex is recruited to telomeric repeats by Sd and E2F1-Dp and modulates telomere length by repressing the transcription of TRs in *Drosophila*. Loss of Sd, Rbf1, or over-expression of E2F1-Dp, stimulate the transcription of TRs, and the effects of Sd, Rbf1, and Cdk8 depletion is dependent on E2F1 and Dp. Furthermore, our CUT&RUN analyses demonstrate direct binding of Cdk8, Dp, and Sd to the repressed telomeric HTT repeats. Collectively, these observations suggest a model for the transcriptional regulation of TRs: the large Mediator complex serves as a direct repressor of TR transcription through Sd and E2F1-Dp during the early S phase (Fig. 5E), while the small Mediator complex is required to activate TR transcription via E2F1-Dp during the G1-S phase transition (Fig. 5F). Remarkably, this model elucidates the precise timing of TR expression, synchronized with the G1-S transition of the host cells, occurring just prior to the elongation of telomeres during the S phase. Our findings highlight a symbiotic relationship between the life cycle of TRs and the cell-cycle machinery of the host cells, resulting in mutual benefits for both.

### Identification of three telomere-specific retrotransposons as *bona fide* transcriptional targets of E2F1-Dp

One unexpected discovery of this work was the identification that three TRs are direct transcriptional targets of E2F1-Dp, the master transcription factor that governs G1-S phase transition. This notion is supported by our following observations (Fig. 4). First, ectopic expression of wild-type E2F1 and Dp is sufficient to stimulate the transcription of all three TRs. Second, transcriptional targets are repressed by Rbf1, and depletion of Rbf1 results in an up-regulation of TR expression. Third, knocking down E2F1 or Dp reduced the transcription of all three TRs. These findings establish a causal link between E2F1-Dp activity and TR transcription. Fourth, the temporal and spatial expression patterns of the TRs in the developing eye and wing imaginal discs correlate well to the cell division cycle and to E2F1 activity. For example, cells within the morphogenetic furrow of eye discs are primarily in G1 phase, while cells posterior to the furrow enter the S phase to initiate the final round of mitosis known as the second mitotic wave (Baker, 2001). We observed an activation of TRs, especially *HeT-A* and TART, in cells located behind the morphogenetic furrow (Fig. S6). Likewise, cells in the anterior compartment of the non-proliferating cell zone in wing discs are arrested in G1 or G2 phases (Johnston and Edgar, 1998). Notably, the expression of TRs in these cells is lower compared to the asynchronously dividing cells outside of this zone (Fig. 1F). These observations provide the biological contexts for the regulation of TR expression by E2F1-Dp. Finally, we identified directly binding of Dp to the TRs in telomeres of XL and 4R (Fig. 5), and the specific E2F1 consensus binding sites can be found in the three TRs. Taken together, these observations suggest the three TRs are *bona fide* transcriptional targets of E2F1-Dp *in vivo*.

### Tight coupling of the telomere-specific retrotransposon life cycle with the cell-cycle machinery of host cells

Considering the crucial role of the RB-E2F network in regulating the G1-S transition, the identification of the three TRs as the direct targets of E2F1 sheds light on a mechanism that links the life cycle of TRs with the cell-cycle machinery of the host cells. The life cycle of TRs is intricately reliant on various components of host cells, for example relying on host RNA polymerase and requisite activating transcription factors, and on translation of the Gag proteins encoded by all three TRs and the reverse transcriptase encoded by *TART* and *TAHRE* (Pardue and DeBaryshe, 2011). Given that the cell-cycle status has profound impacts on diverse cellular processes, it is perhaps not surprising that the life cycle of TRs, which heavily relies on various components of host cells, must adapt to the cell-cycle status of the host cells to ensure the success of both TRs and host cells.

While most eukaryotes resolve the “end replication problem” using telomerase to add telomere repeats to chromosome ends, Dipteran insects have evolved an alternative using TR retrotransposition. A common feature of these two mechanistic solutions is the coupling of telomere elongation with telomere replication during S phase. HeT-A mRNA is most abundant during S phase (George and Pardue, 2003; Walter and Biessmann, 2004), and Orf1p (of *HeT-A*) is predominantly detected at the G1/S boundary and in early S phase, but not in M phase (Zhang et al., 2014). Thus, our finding that E2F1-Dp directly regulates the transcription of all three TRs exactly explains the mechanism of cell cycle regulation of telomere addition. These findings reveal a robust coupling between the life cycle of TRs and the cell-cycle machinery in host cells, ensuring efficient maintenance of telomere homeostasis and genomic stability across multiple cell cycles. Therefore, this intricately intertwined strategy is mutually beneficial for both TRs and host cells.

### Dual role of the Mediator complex in regulating telomere-specific retrotransposon transcription

The CDK8 module binds to the small Mediator complex and forms the large Mediator complex, which generally represses Pol II-dependent transcription (Allen and Taatjes, 2015). Depletion of multiple subunits of the large Mediator complex leads to similar up-regulation of TR transcription, indicating that the large Mediator complex negatively affects the transcription of TRs. The Mediator complex only binds to DNA through interacting with specific transcription factors. In *Drosophila*, five Mediator subunits, Med1, Med15, Med19, Med23, and Med31, bind the Yki transcription factor (Oh et al., 2013), similar to human Mediator subunits Med12, Med14, Med23, and Med24 bind to YAP in bile duct carcinoma cells (Galli et al., 2015). This appears to be a general mechanism as ChIP-seq analyses show that 87% of YAP-binding sites overlap with Med1 binding sites (Galli et al., 2015). Moreover, the Yki homolog TAZ has been shown to interact with MED15 in human embryonic stem cells (Varelas et al., 2008). Depleting either *sd* or *yki* leads to up-regulation of TR transcription. Therefore, a simple model is that the recruitment of the Mediator complex to the HTT repeats is facilitated through the direct binding of Sd-Yki to DNA.

Additionally, we have observed that the knockdown of *Rbf1*, which represses E2F1-Dp activity, leads to a strong up-regulation of TR expression. Remarkably, depletion of the Mediator complex has a considerably weaker effect on TR transcription compared to depleting *Rbf1*. We postulate that E2F1-dependent transactivation of TRs relies on the small Mediator complex, which implies that knocking down some shared Mediator subunits would partially disrupt both TR repression by the large Mediator complex and E2F1-dependent TR transactivation by the small Mediator complex. We previously showed that two subunits of the Mediator complex, Cdk8 and CycC, negatively regulate E2F1-dependent transcription (Morris et al., 2008), perhaps through E2F1 phosphorylation by Cdk8 (Morris et al., 2008; Zhao et al., 2013).

Our results favor the model in which the small Mediator complex is involved in E2F1-dependent transcription of TRs. To fully elucidate this mechanism, further studies aimed at identifying the specific Mediator subunits directly interacting with E2F1 or Dp will be necessary. It is conceivable that the Mediator complex plays a dual role in regulating TR transcription. During the early G1 phase, the large Mediator complex represses TR transcription, primarily via Sd-Yki (Fig. 5E). In contrast, during the G1-S phase transition and in early S phase, the small Mediator complex becomes essential for activating TR transcription, primarily through E2F1-Dp (Fig. 5F).

### Context-dependent crosstalk between Sd-Yki and RBF1-E2F1-Dp complexes

Sd/TEAD serves as a critical transcription factor downstream of the Hippo signaling pathway, playing conserved roles in controlling organ size, and frequently undergoing dysregulation in human cancers (Pan, 2010; Yu et al., 2015). The interplay between the Rb-E2F1 regulatory network and Hippo signaling helps to regulate cell-cycle exit and apoptosis (Nicolay et al., 2011; Zhang et al., 2017). In the eye imaginal discs, E2F1 synergizes with Yki-Sd to activate the expression of shared target genes, leading to excessive cell proliferation and inappropriate tissue growth (Nicolay et al., 2011). Interestingly, E2F1 also regulates apoptosis and organ size in *Drosophila* wing imaginal discs by repressing the expression of Yki targets, such as *Diap1*, *expanded*, and *bantam* (Zhang et al., 2017). The underlying mechanism involves the competition between E2F1 and Yki for binding to Sd, resulting in the formation of the E2F1-Sd repressor complex, and Rbf1 modulates this process by reducing the interaction between E2F1 and Sd (Zhang et al., 2017). This molecular mechanism is conserved in humans (Zhang et al., 2017). These data suggest that E2F1 interferes with the binding of Yki/YAP to Sd/TEAD, thereby suppressing the expression of Yki-YAP target genes (Zhang et al., 2017), however the interplay between the RBF-E2F1 and Hippo pathways is further complicated by the fact that dE2f1 is a transcriptional target of Sd-Yki (Goulev et al., 2008; Zhang et al., 2017). This potential feedback loop likely regulates the activities of both pathways. If so, it is also likely that the intricate interplay between RBF-E2F1-Dp and Sd-Yki is probably shaped by the specific promoter structures of shared target genes and the distinct contexts of the biological processes in which they are involved. Supporting this idea, Yki-Sd and E2F1 can activate common targe genes such as *Dachs*, *Dp*, and *PCNA* in eye discs (Nicolay et al., 2011).

In this study, we observed that RBF-E2F-Dp and Sd-Yki play important roles in regulating TR transcription and telomere homeostasis. Our results suggest that Sd and Yki negatively regulate TR expression, whereas E2F1 and Dp are required for TR transcription. We propose that this protein complex is responsive to signaling from growth factors and other extracellular stimuli that regulate cell proliferation. It would be intriguing to explore whether additional components of the Hippo signaling pathway, as well as other signaling pathways, also regulate the transcription of TRs through Mediator. Studies of *Drosophila* telomeres – despite them being “non-canonical” – have long revealed features of chromosome biology, and how retrotransposons and host genomes coevolve (Pardue and DeBaryshe, 2011). Our work unravels how the life cycle of TRs are closely coupled with the cell-cycle machinery of the host cells, revealing how the transcription of TRs are tied to the G1-S phase transition, ahead of retrotransposition happening during DNA replication in S phase.

## Acknowledgement

We thank Jian-Quan Ni for assistance in generating the *Cdk8^ΔmCherry^* allele, Jasmine Sun for technical support, and Henri-Marc Bourbon and Lori Walrath for sharing fly strains and reagents. We appreciate the insightful comments by Nick Dyson and Keith Maggert on the manuscript. We also thank the Bloomington *Drosophila* Stock Center (NIH Grant P40OD018537) for their provision of many fly stocks. Z.L. is partially supported by the National Institute of Health grants R01CA261258 and P20GM121288. This research was supported by a grant from the National Institute of Health (GM133011 to J.-Y.J.).

## Materials and Methods

### Fly stocks and maintenance

Flies were maintained at 25°C on a standard medium consisting of cornmeal, molasses, and yeast. The *w^1118^ Drosophila* strain was used as the control group. Null alleles of *dCdk8* (*FRT80B dCdk8^K185^/TM3 Sb*) and *CycC* (*FRT82B CycC^Y5^/TM3 Sb*) were provided by Henri-Marc Bourbon. The *118E-15* strain used for TPE assays were obtained from Lori Walrath. The specific *Drosophila* strains and their genotypes are listed in Table S1.

### Analyses of polytene chromosomes

To analyze larval polytene chromosomes, we followed the established procedure as described previously (Xie et al., 2015).

### Generation of the *dCdk8^ΔmCherry^* and *dCdk8^EGFP^*alleles using the CRISPR-Cas9 technique

To generate the *dCdk8^ΔmCherry^* allele, we used CRISPR/Cas9-mediated homology-directed repair approach (Ren et al., 2014). We designed two guide RNAs (*dCdk8-sgRNA1*: AACACAGCCTTAACCAGGGA, *dCdk8-sgRNA2*: TCGTTGAAATATCTTTCCGA) using the online CRISPR design tool available at http://www.flyrnai.org/crispr/. These sgRNAs were designed to create double strand breaks near the transcription start and termination sites of the *dCdk8* gene. Next, we inserted the sgRNAs into the *U6b*-sgRNA-short vector following the procedure as previously described (Ren et al., 2013). Meanwhile, we constructed the *dCdk8-4XP3-mCherry* donor vector as previously described (Ren et al., 2014). For the injection process, we combined the plasmid mix as follows: *dCdk8-sgRNA1* at 75 ng/μl, *dCdk8-sgRNA2* at 75 ng/μl, and *dCdk8-4XP3-mCherry* at 100 ng/μl. These plasmids were injected into embryos obtained from *P{nos-Cas9}attP40* flies, resulting in the generation of the *dCdk8^ΔmCherry^* mutants. To identify successful *dCdk8^ΔmCherry^* mutants, we screened for the expression of mCherry in the eyes using a fluorescent microscope, followed by genotyping PCR and Sanger sequencing to confirm the desired mutations.

To create the *dCdk8^EGFP^* allele, we used the CRISPR Optimal Target Finder tool available at http://targetfinder.flycrispr.neuro.brown.edu to design two sgRNAs. These sgRNAs were positioned near the transcription start site and transcription termination site to replace the entire *dCdk8* gene. We then cloned those two sgRNAs into the *pCFD3-dU6:3gRNA* vector following the protocol outlined previously (Port et al., 2014), and the primer redesign was facilitated using the NEB Builder tool, with the primer sequences provided in Table S2 for reference. For the assembly of the doner constructs, we used the NEBuilder HiFi DNA Assembly Cloning Kit (NEB #E5520). This involved integrating both upstream and downstream homology arms, each spanning 1000bp, along with the last intron-exon regions and the EGFP coding fragment, into the pGEM-T vector (Promega A1360). The *dCdk8-EGFP* rescue construct, as described previously (Xie et al., 2015), served as the PCR template for amplifying the EGFP-fused *dCdk8* gene region. The injection of embryos with the donor vector and the two sgRNA constructs within the pCFD3 vector was conducted by Rainbow Transgenic Flies (https://www.rainbowgene.com/). Transgenic flies carrying the EGFP tag at the C-terminus of dCdk8 were identified through classical fly genetics methods. Validation was subsequently carried out by performing on extracted genomic DNA and confirming the results though sequencing.

### Western Blot Analysis

Western blot analysis was conducted using the same protocol as previously described (Xie et al., 2015). We used the following antibodies: anti-dCdk8 (polyclonal antibody from guinea pig, diluted at 1:1000; kindly provided by Henri-Marc Bourbon) (Gobert et al., 2010), and anti-actin monoclonal antibody (diluted at 1:4,000, Thermo Scientific).

### RNA preparation, quantitative Reverse-Transcription-PCR (qRT-PCR), and RNA-seq analysis

To isolate total RNA from third instar wandering larvae or dissected larval central nervous system, we used the Trizol reagent (Invitrogen). The isolated RNAs were quantified, treated with DNase I to eliminate any genomic DNA contamination, and reverse transcribed using the High-Capacity cDNA Reverse Transcription kit from Applied Biosystems. In our quantitative PCR (qPCR) analysis, we used SYBR Green from Applied Biosystems; the qPCR primers are listed in Table S2. *Rp49* was used as the internal control, using the same primer pair as previously described (Xie et al., 2015). For RNA-seq analysis, we followed the established protocol as described previously (Li et al., 2022). The RNA-seq data related to this study have been deposited in NCBI’s Gene Expression Omnibus and are accessible through the GEO Series accession number GSE217104.

### DNA extraction and qPCR

For DNA extraction, five wandering stage larvae were collected and homogenized in 500µl of squishing buffer, consisting of 0.1 M Tris-HCl (pH 9.0), 0.1 M EDTA, and 1% SDS. The homogenate was incubated at 70°C for 30 minutes. Next, 70 µl of 8.0 M potassium acetate was added to the homogenate, and samples were left on ice for an additional 30 minutes. The mixture was centrifuged at 12,000 rpm at 4°C for 15 minutes, and the supernatant was transferred into an Eppendorf tube. To precipitate the DNA, 0.5 volumes of isopropanol were added to the supernatant. The mixture was then centrifuged again at 12,000rpm for 5 minutes to pellet the DNA, which was then washed with 1.0 ml of 70% ethanol and was resuspended in 50µl distilled water. For qPCR analysis, we used SYBR Green (Applied Biosystems) and the same primers used in the qRT-PCR analyses (Table S2).

### The HCR RNA-FISH assay

We followed the protocol for the multiplexed in situ HCR as previously described (Li et al., 2022). The following probe sets and amplifiers were obtained from Molecular Instruments: the B1-Alexa Fluor 488 amplifiers were used in conjunction with the probe set designed for *HeT-A* (lot number PRO329); the B2-Alexa Fluor 594 amplifiers were used with the probe sets designed for *TART* (lot number PRO330); and the B3-Alexa Fluor 647 amplifiers were applied alongside with the probe sets designed for *TAHRE* (lot number PRQ334). Confocal images were acquired using a Zeiss LSM900 confocal microscope system and subsequently processed using Adobe Photoshop 2021.

### Identification of the genome-wide Cdk8-, Dp-, and Sd-binding sites through CUT&RUN sequencing

We performed the CUT&RUN assay following the same protocol outlined previously (Liu et al., 2023). The analysis of the CUT&RUN sequencing data was conducted using *Drosophila* genome (assembly Release r5.9) as the reference. Upon acceptance of this work, we plan to deposit the sequencing results from the CUT&RUN assay in NCBI’s Gene Expression Omnibus for public access. To visualize the gene tracks, we used the Integrative Genomics Viewer (IGV 2.14.1) browser, setting the y-axis to autoscale for optimal presentation and clarity.

### Statistical analyses

For each genotype analyzed in this study, we performed a minimum of three independent biological replicates. P-values were calculated using Microsoft Excel, and standard deviation is represented by error bars in the figures. Significance levels, determined using one-tailed unpaired *t*-tests, were denoted as follows: * p<0.05; ** p<0.01; *** p<0.001.

---

**Fig. S1. Impact of *Cdk8* and *CycC* mutation on telomere length and TPE.** (A-D) Polytene chromosome staining was performed to assess telomere ends in heterozygous mutant larvae, including *Cdk8^K185^*/+ (B), *CycC^Y5^*/+ (C), and *Cdk8 ^ΔmCherry^*/+ (D). Arrows highlight telomeres, note the uneven telomere ends observed in polytene chromosomes from both *Cdk8* and *CycC* heterozygous mutants compared to the control (A; *w^1118^*/+). (E-I) Dominant enhancement of TPE was examined through the mottled eye color phenotype in male adult flies (*118E-15*). The genotypes included: (E) *w^1118^*/Y; *+; +; 118E-15/+*; (F) +/Y; *+; Cdk8^K185^/+; 118E-15/+*; (G) +/Y; *+; CycC^Y5^/+; 118E-15/+*; (H) +/Y; *+; Cdk8^K185^ CycC^Y5^/+; 118E-15/+*; and (I) +/Y; *+; Cdk8 ^ΔmCherry^/+; 118E-15/+.* (J) Cdk8 protein levels were assessed in *w^1118^* and *Cdk8 ^ΔmCherry^* mutant larvae through Western Blot analysis. (K/L) Detection of TR transcripts in salivary gland cells using the HCR RNA-FISH assay.

**Fig. S2. Characterization of the *dMed7^MI10755^* mutant allele.** (A) The genomic region of the *Med7* locus is depicted, with a green arrow indicating the insertion site of *dMed7^MI10755^*. (B) The *dMed7^MI10755^* allele was validated through PCR using the primers (*dMed7F* and *dMed7R*) indicated in (A). No PCR products were detected in genomic DNA from *dMed7^MI10755^* homozygous mutant larvae due to the transposon insertion. (C) Homozygous mutants of *dMed7^MI10755^*exhibit developmental arrest at the third instar larval stage. (D) Analysis using qRT-PCR revealed an upregulation of *HeT-A* and *TART* transcripts in *dMed7^MI10755^* mutant larvae. (E) Dominant enhancement of TPE by *dMed7^MI10755^* allele.

**Fig. S3. Effect of Sd on TR expression and telomere homeostasis.** (A/B) Polytene chromosome staining was performed to assess telomere ends in polytene chromosomes from the control *w^1118^*/+ (A) and *sd^1^/+* (B) heterozygous mutant larvae. Arrows highlight the telomeres, note the uneven telomere ends in in *sd^1^/+* heterozygous mutants. (C/D) Dominant enhancement of TPE of *118E-15* mottled eye color phenotype in female adult flies. Genotypes included: (C) *w^1118^*; *+; +; 118E-15/+*; and (D) *sd^1^*/*w^1118^; +; +; 118E-15/+*. (E/F) The expression of *HeT-A* (green), *TART* (orange), and *TAHRE* (magenta) transcripts was detected using the HCR RNA-FISH assay in ovary tissue. Notably, (F) shows elevated TR expression in nurse cells in egg chambers from *sd^1^* mutant females compared to the control (E).

**Fig. S4. Effect of E2F1-Dp and Rbf1 on TR expression in wing imaginal discs as detected by the HCR RNA-FISH assay.** Genotypes: (A, control) *en-Gal4/+; UAS-BFP/+*; (B) *en-Gal4/+; UAS-E2f1^+^ UAS-Dp^+^/UAS-BFP*; and (C) *en-Gal4/+; UAS-Rbf1^RNAi^/UAS-BFP*. Notably, in the posterior compartment of the wing discs, both overexpression of E2F1-Dp and depletion of Rbf1 significantly increase the mRNA levels of all three TRs (*HeT-A* in green; *TART* in orange; and *TAHRE* in magenta).

**Fig. S5.** Genomic tracks are presented, highlighting binding peaks for Cdk8 (in red), Dp (in green), and Sd (in blue) at the telomeric regions of four different chromosomes: (A) Telomere of 2R (with the chromosome end on the *right*), (B) Telomere of 4L (with the chromosome end on the *left*), (C) Telomere of 3L, and (D) Telomere of 3R. Note that these telomeric regions have limited annotation in the available *Drosophila* genome (assembly Release r5.9).

**Fig. S6.** mRNA transcripts of *HeT-A* (green, B), *TART* (orange, C), and *TAHRE* (magenta, D) detected using the HCR RNA-FISH assay in wild-type wing discs during the late third instar larval stage. The arrows highlight the expression of TRs in cells behind the second mitotic wave.

**Table S1.** List of the *Drosophila* strains used in this study.

**Table S2.** Primers used in qPCR assays and the generation of the *dCdk8^EGFP^* strain using the CRISPR-Cas9 technique.

