## Supplementary figures and images for "Transcriptional coupling of telomeric retrotransposons with the cell cycle"

### Supplementary Figures 1-6

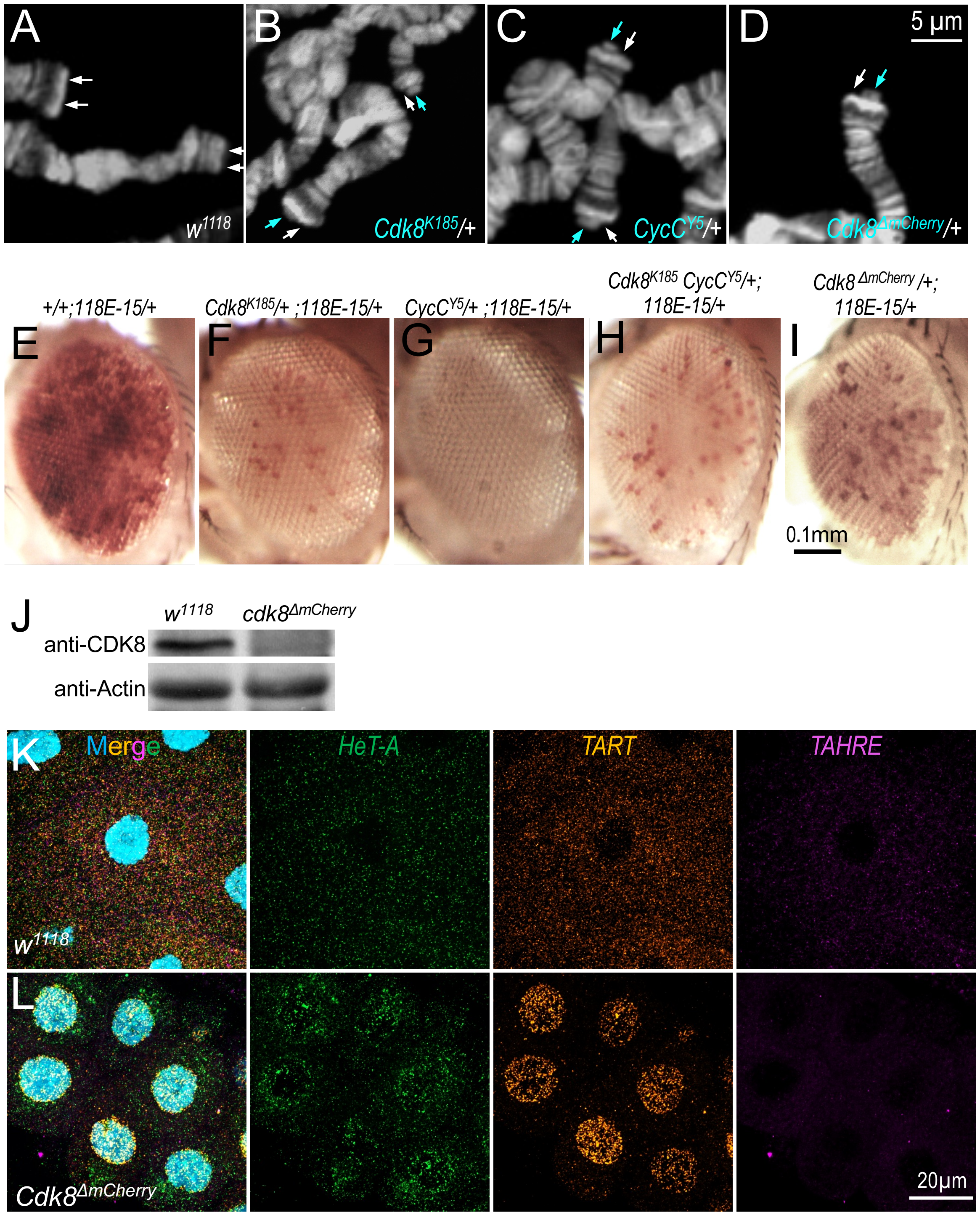

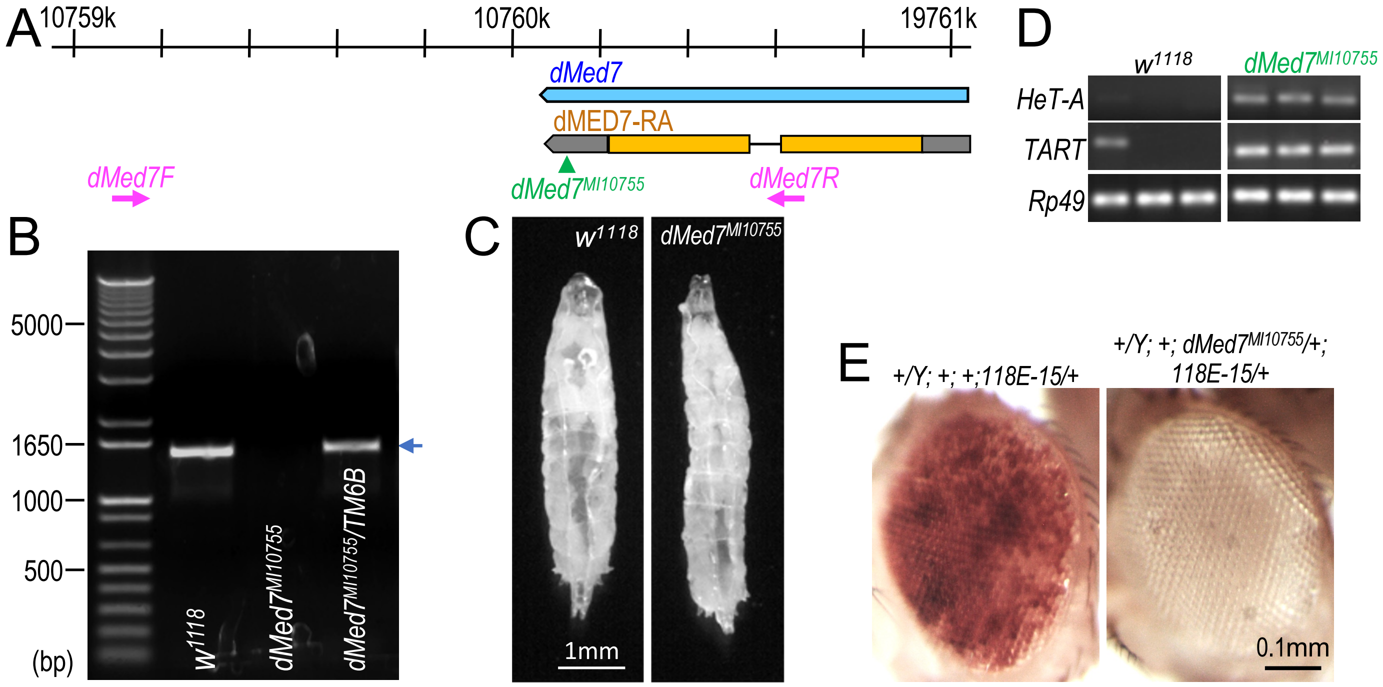

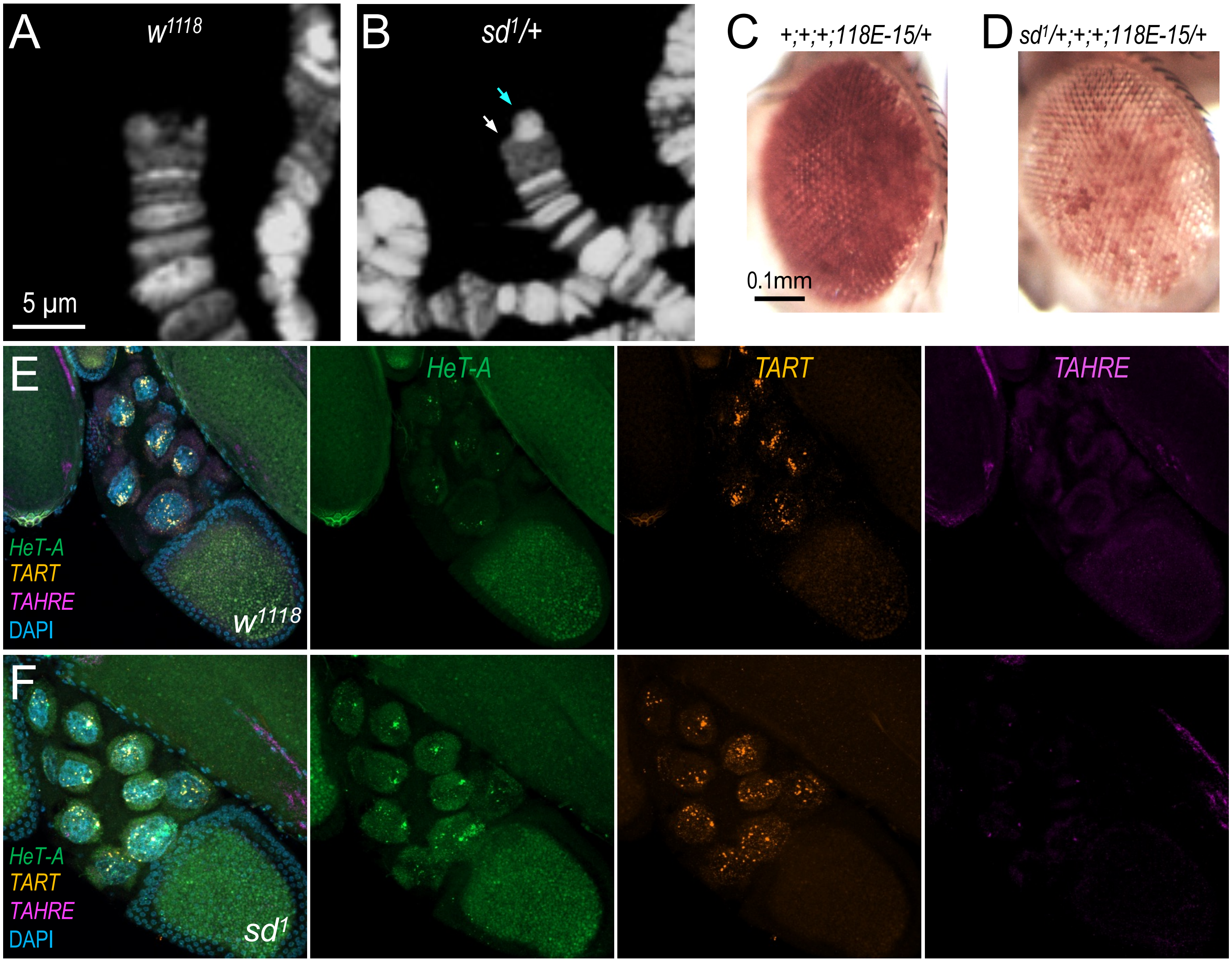

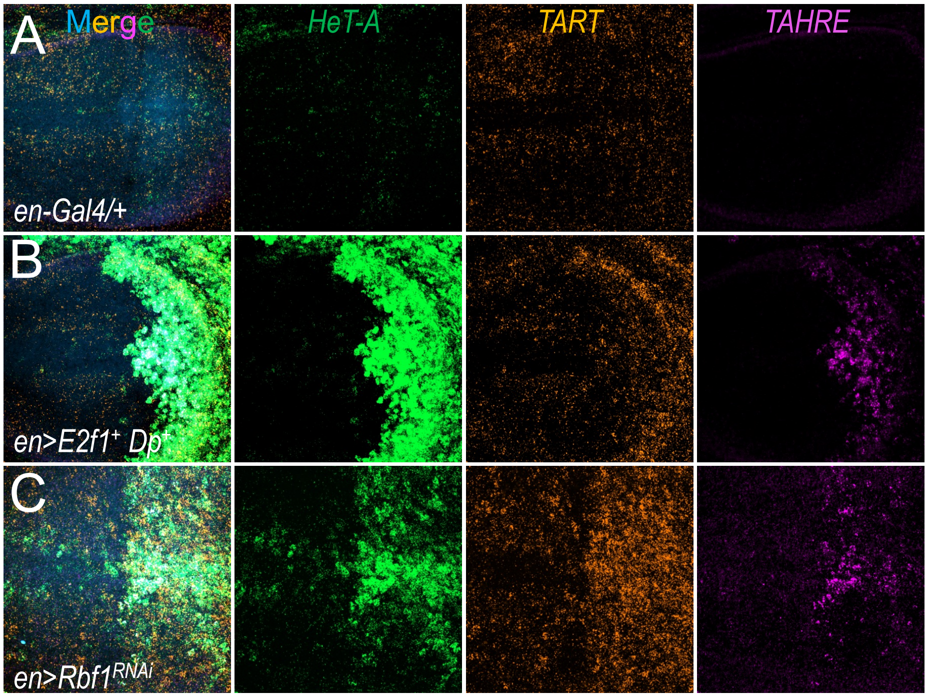


**Fig. S4**

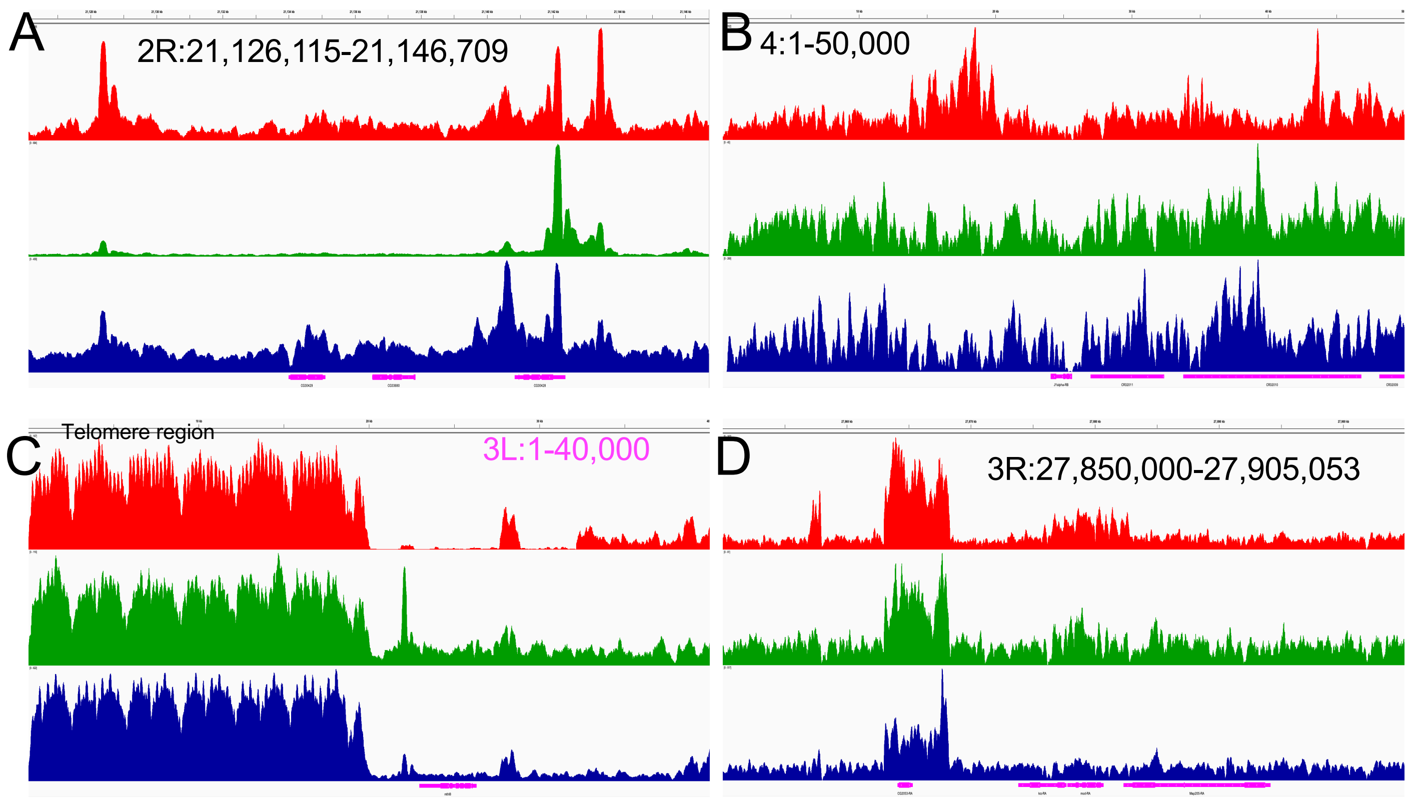

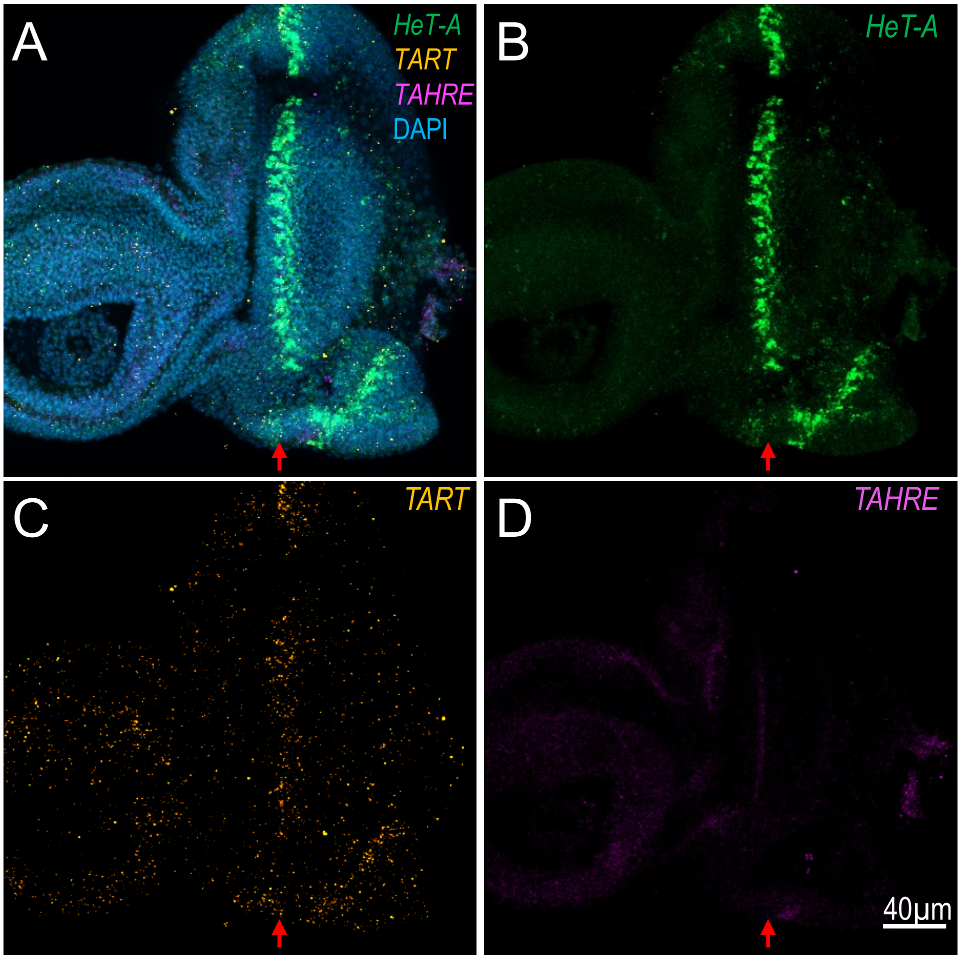


**Fig. S6**
